# ClusterApp to visualize, organize, and navigate metabolomics data

**DOI:** 10.1101/2025.02.12.637912

**Authors:** Vinicius Hansel, Pothuvilage Karunarathne, Tiago Cabral Borelli, Robert Quinn, Ricardo R. da Silva

**Affiliations:** Department of Biomolecular Sciences, Computational Chemical Biology Laboratory, School of Pharmaceutical Sciences of Ribeirão Preto, University of São Paulo, Ribeirão Preto, Brazil; Department of Biochemistry and Molecular Biology, Michigan State University, East Lansing, USA

**Author notes:** Corresponding authors RR da Silva -, R Quinn. Contributed equally (Vinicius Hansel, Pothuvilage Karunarathne).

## Abstract

**Background:** Clustering analysis is a foundational step in exploratory data analysis workflows, with dimensionality reduction methods commonly used to visualize multidimensional data in lower-dimensional spaces and infer sample clustering. Principal Component Analysis (PCA) is widely applied in metabolomics but is often suboptimal for clustering visualization. Metabolomics data often require specialized manipulations such as blank removal, quality control adjustments, and data transformations that demand efficient visualization tools. However, the lack of user-friendly tools for clustering without computational expertise presents a challenge for metabolomics researchers. ClusterApp addresses this gap as a web application that performs Principal Coordinate Analysis (PCoA), expanding clustering alternatives in metabolomics. Built on a QIIME 2 Docker image, it enables PCoA computation and Emperor plot visualization. The app supports data input from GNPS, GNPS2, or user-provided spreadsheets. Freely available, ClusterApp can be locally installed as a Docker image or integrated into Jupyter notebooks, offering accessibility and flexibility to diverse users.

**Results:** To demonstrate the data preprocessing techniques available in ClusterApp, we analyzed two Liquid Chromatography coupled to Tandem Mass Spectrometry (LC-MS/MS) metabolomics datasets: one exploring metabolomic differences in mouse tissue samples and another investigating coral life history stages. Among the dissimilarity measures available, the Bray-Curtis measure effectively highlighted key metabolomic variations and patterns across both datasets. Targeted filtering significantly enhanced data reliability by retaining biologically relevant features, 10,617 in the coral dataset and 7,341 in the mouse dataset while eliminating noise. The combination of Total Ion Current (TIC) normalization and auto-scaling improved clustering resolution, revealing distinct separations in tissue types and life stages. ClusterApp’s flexible features, such as customizable blank removal and group selection, provided tailored analyses, enhancing visualization and interpretation of metabolomic profiles.

**Conclusion:** ClusterApp addresses the need for accessible, dynamic tools for exploratory data analysis in metabolomics. By coupling data transformation capabilities with PCoA on multiple dissimilarity matrices, it provides a versatile solution for clustering analysis. Its web interface and Docker-based deployment offer flexibility, accommodating a wide range of use cases through graphical or programmatic interactions. ClusterApp empowers researchers to uncover meaningful patterns and relationships in metabolomics data without requiring cumbersome data manipulation or advanced bioinformatics expertise.

## Background

Modern analytical techniques, such as mass spectrometry and DNA sequencing, generate increasingly large volumes of data [1]. Mining this data for biological insights is challenging, requiring tools that can help compress the multivariate information into digestible and interpretable visualizations. Many dimensionality reduction methods have been around for decades, including Principal Component Analysis (PCA) and Multidimensional Scaling (MDS) [2]. These methods are often used to visualize multivariate data by reducing its complexity while preserving as much variation as possible, enabling researchers to identify patterns, clusters, and relationships within high-dimensional datasets [3]. Most often, it is computed through statistical software, such as R, MATLAB, or similar platforms requiring at least some knowledge of computer programming. Other sources of computing are online tools that require the creation of an account and asynchronous processing in a complex computational environment, for example, with GNPS or XCMSOnline [4,5]. Even with extensive experience in such computation and data interpretation, it is still a cumbersome process to organize, visualize, and analyze this multivariate data regardless of the software platform. This is especially true in the case of metabolomics data, a multivariate data type that is similar to sequencing-based omics, such as transcriptomics, metagenomics, and others, but unique in some aspects of its data structure and handling requirements [6,7,8]. One aspect of metabolomics that creates challenges in visualization in multivariate space is the need to check quality control samples, blank samples, and other quality assurance measurements that can help navigate contaminants in analytical instruments to ensure high data quality. While not exclusive to metabolomics, this need to check, clean, and visualize data in multivariate space in an iterative process creates the need for tools that can enable these manipulations quickly and easily to visually identify any data problems and structure of interest uncovering meaningful patterns and relationships. This is especially critical in untargeted metabolomics, where data variability can mask meaningful trends if not properly managed, and addressing this variability is essential for handling the large volumes of data typical in untargeted metabolomics, ultimately facilitating clearer interpretation and deeper insights [8].

To enhance the flexibility of metabolomics data analysis, it is essential to offer tools that support a wide array of data processing and visualization techniques. This includes capabilities for normalization, scaling, and dissimilarity measure selection, which are critical for obtaining accurate insights from complex datasets. The majority of metabolomics studies use only PCA for exploratory data analysis which is the method implemented on the most popular open-source tools, such as MetaboAnalyst [9] and XCMSOnline. Although tools like GNPS [4] implement PCoA analysis as standard results for some of its workflows, the platform does not allow the user input option to normalization, scaling, and dissimilarity measure selection. For the microbiome community, it has been shown that the ability to detect clustering signatures for longitudinal or treatments vs control studies is dependent on the choice of dissimilarity measure [10]. Therefore, we aimed to create an easy to use web app, enabling the user to quickly perform multiple transformations in their data and promptly visualize the impacts on a lower dimensional representation.

## Implementation

Principal Component Analysis (PCA) is a linear dimensionality reduction that transforms the original data to a new coordinate system created by a scalar projection, the principal components (PCs), which are ordered by the variance explained, with the greatest variance on the first PC, and so on [11]. For an *n x p* data matrix, X, where the *n* rows represent the different experimental samples and the *p* columns represent the measured variables (*e*.*g*., metabolites, proteins, genes, etc.), the goal is to find a linear combination of the original variables that have maximum variance, and therefore, captures as much of the original data variability as possible.

Classical Multidimensional Scaling - MDS (also known as Principal Coordinate Analysis (PCoA)) tries to find a set of pairwise distances between *n* samples given a *n*×*n* matrix **D** [12]. This task is accomplished by finding a low-dimensional embedding of data points in ℝ^k^, where k represents the number of axes on the lower dimensions space, such that Euclidean distances between them approximate the given distances. The approximation is obtained by minimizing a cost function called “stress”. PCoA can be computed on non-euclidean distances, which can be more biologically meaningful for a wide range of studies [13]. It can be shown that PCoA performed on an Euclidean distance matrix is equivalent to PCA [14]. Therefore, PCoA offers greater flexibility for exploratory data analysis when searching for new patterns in experimental data.

Web applications are essential for many research fields, enabling scientists with little or no knowledge of computer programming or command line interface tools to perform complex analysis through graphical web interfaces. In the field of metabolomics, the previously mentioned web platforms GNPS, MetaboAnalyst, and XCMSOnline have thousands of citations, being used by thousands of users around the world. Besides lacking flexible implementations for performing PCoA analysis, these platforms are also difficult to implement and customize locally. To overcome these limitations and provide a PCoA visualization approach agnostic to the metabolome data processing platform, ClusterApp is implemented in the microframework Flask, which allows simplicity and flexibility for data analysis (Figure 1). The implementation features a direct connection to GNPS, enabling the retrieval of data from different GNPS and GNPS2 workflows, in addition to user-provided spreadsheets. The web interface also features a data formatting functionality, allowing data analysis and reformatting cycles. All these functionalities are explained in video and text formats. The application extends a QIIME 2 Docker image, allowing seamless local installation, as well as custom analysis using example Jupyter notebooks. ClusterApp performs the most commonly used normalization (Total ion current/TIC and Probabilistic Quotient Normalization/PQN) as well as the most commonly used scaling (autoscaling and pareto) methods reported in metabolomics literature [15,16]. Additionally, ClusterApp uses the SciPy distance module to allow users to calculate various dissimilarity metrics [17]. Finally, ClusterApp uses QIIME 2’s API to perform PCoA calculations and Emperor to create interactive visualizations [18,19].

**Figure 1.**
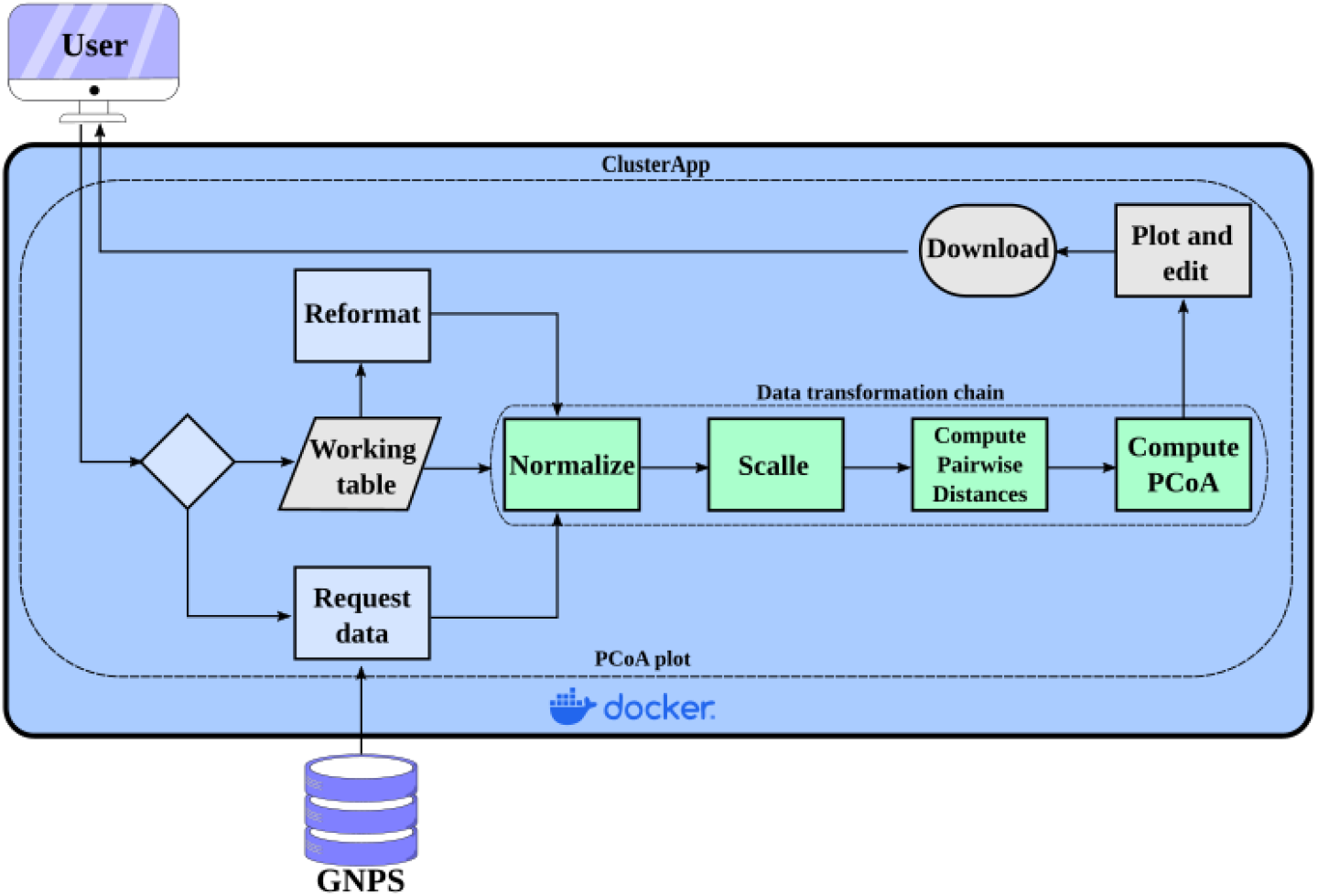
ClusterApp web application model. The user can retrieve data from GNPS or GNPS2, or provide a spreadsheet in a designed format, described on the tool’s ‘Usage’ page. The PCoA results are displayed on the browser and can also be downloaded for easy sharing.

## Results and discussion

### Enhanced clustering capabilities through PCoA in ClusterApp

The current ClusterApp implementation allows PCoA analysis (Figure 2a), which expands the clustering recognition beyond PCA (Figure 2b), by allowing the selection of multiple different dissimilarity metrics. PCoA is particularly advantageous for analyzing complex datasets where flexibility in choosing distance metrics is essential. Unlike PCA, which relies solely on Euclidean distances, PCoA can compute dissimilarities using a wide variety of metrics [1]. ClusterApp further enhances the utility of PCoA by offering users a selection of 20 dissimilarity measures, including Euclidean, Bray-Curtis, and Jaccard, allowing for tailored analyses based on dataset characteristics. This flexibility makes PCoA analysis in ClusterApp a powerful tool for identifying and visualizing patterns in metabolomic data.

**Figure 2.**
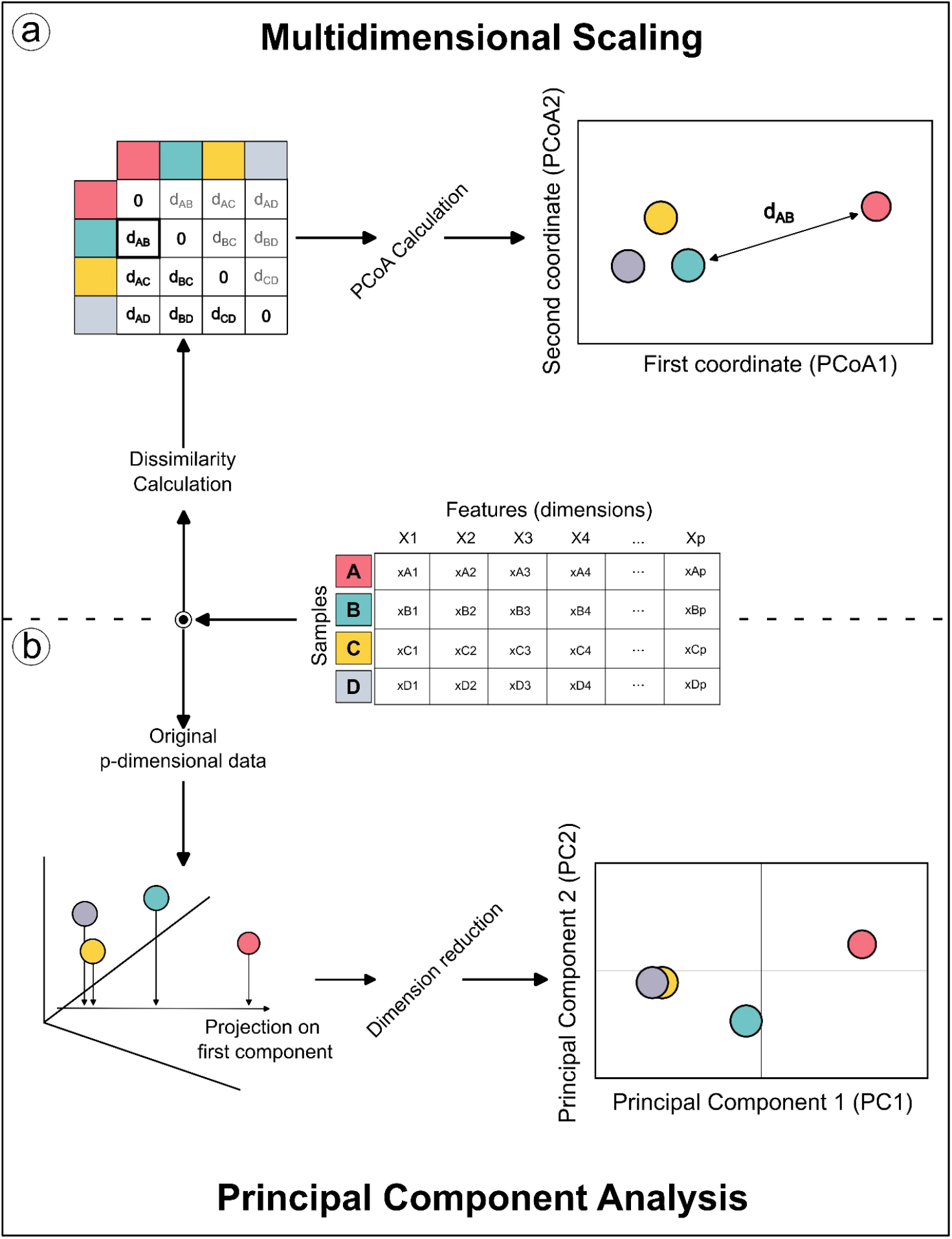
Schematic representation of differences between PCoA and PCA. A) In PCoA computation, a dissimilarity matrix, which can vary according to one’s choice, is calculated from an original dataset, and the samples are represented in the new coordinate system. B) For PCA, the original data points are used to calculate the projection onto a new coordinate system.

### Streamlined data cleaning and preprocessing with ClusterApp

Data preprocessing is essential for ensuring reliable and reproducible metabolomic analysis, addressing challenges such as background noise, outliers, and data inconsistencies. Efficient data cleaning and preprocessing are crucial for identifying patterns, trends, and anomalies, which accelerate insights and guide informed decision-making. ClusterApp simplifies this process with feature filtering based on blank sample intensity, normalization, and scaling, enhancing data comparability and clustering.

To demonstrate the various data preprocessing techniques available in ClusterApp, we utilized two metabolomics datasets as case studies (Supplementary Datasets 1 and 2): one examining metabolomic differences in mouse tissue samples (Figure 3A) and another exploring coral life history stages (Figure 3B), both generated via LC-MS/MS. We employed various dissimilarity measures available in ClusterApp, with the Bray-Curtis dissimilarity measure effectively highlighting metabolomic variation and revealing key and biologically anticipated metabolomic patterns in both datasets.

**Figure 3.**
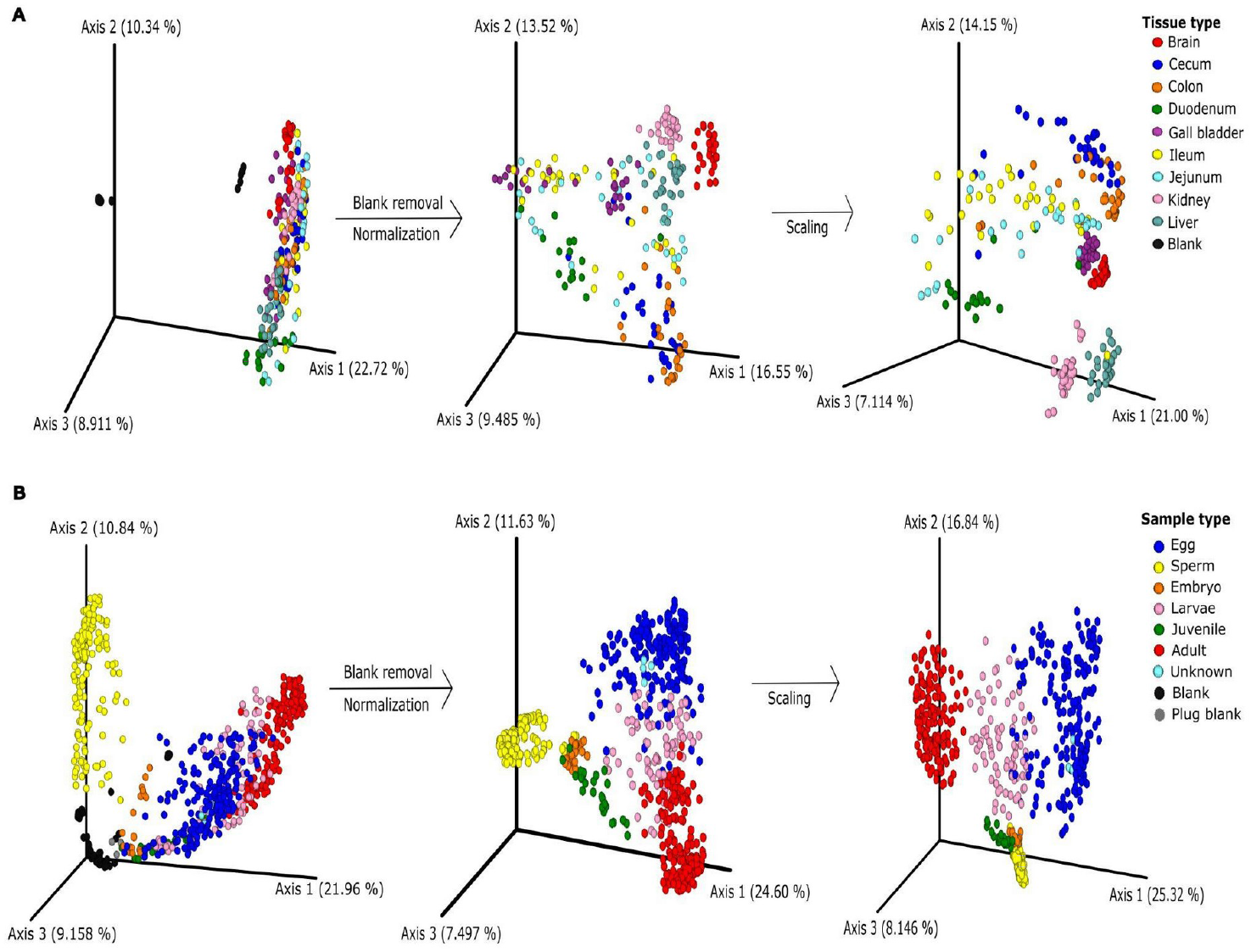
Application of data pre-processing techniques in ClusterApp. PCoA plots demonstrating cluster analysis of metabolomics data before and after applying blank removal, normalization, and scaling in A) different tissue samples in mice and B) life history stages of coral samples. The blank removal was carried out using a threshold of 2× the average intensity of blanks across at least 5% of non-blank samples. The data were normalized using TIC (Total Ion Current) and scaled using the auto-scaling options available in ClusterApp. Bray-curtis dissimilarity measure was used to generate the PCoA plots and the plots are color-coded by tissue or sample type.

### Feature filtering based on blank sample intensity

Blank removal is an optional feature in ClusterApp, designed to help users eliminate background features originating from extraction solvents, environmental contaminants, and instrument-related noise. These signals can obscure low-intensity analytes, complicating accurate detection and quantification. ClusterApp employs a filtering approach that retains features with intensities exceeding a user-defined threshold relative to the average blank sample intensity across a specified proportion of non-blank samples. For example, features with intensities ≥3× the average blank intensity in 90% of non-blank samples can be retained. As blank sample types and positions on an analytical acquisition sequence may vary depending on the experimental design [20], this feature should be used with care and the user should take note of the number of features being removed, which is informed by the app, in each combination of parameters used to filter the data.

In our case study, we applied ClusterApp’s intensity-based blank removal feature to analyze two LC-MS/MS datasets: metabolomic differences in mouse tissue samples (Figure 3A) and coral life history stages (Figure 3B). First, we applied a threshold to keep features whose average intensity in at least 50% of the non-blank samples should be higher than 30% of the average intensity in samples tagged blank. As expected we found only 546 features (5% of total features) meeting the criteria. Due to the sparse nature of feature tables, it is common to find features only in a small subset of samples, therefore the average intensity would not be representative of a skewed distribution. Such a filter would be even more strict if the experimental groups are expected to vary greatly or if the blank samples are not representative of the sample set. One possibility would be to require that at least a subset of non-blank samples, say 5% are at least 2x (200%) more intense than blank samples. With these parameters, we recovered 6,340 features (58%) in the mouse tissue dataset. Similarly, for the coral dataset, 5,246 features (70%) met the criteria. This conservative approach would allow some experimental groups or samples to have features independently represented in the final dataset.

In both datasets, blank samples clustered distinctly from the rest of the data, a trend readily visible in ClusterApp’s interface. This initial visualization highlights the critical role of blank evaluation in reducing background noise and enhancing data precision. The intensity-based blank removal approach retains features that exceed a user-defined intensity threshold relative to the average blank sample intensity in a specified proportion of non-blank samples, ensuring that only biologically relevant signals are preserved. By minimizing noise while retaining meaningful features, this targeted filtering method enhances data reliability. The customizable nature of ClusterApp’s blank removal feature allows users to adjust thresholds. This flexibility ensures that the analysis can be tailored to the unique characteristics and research goals of each dataset, providing a robust and adaptable solution for metabolomic studies.

### Normalization

ClusterApp offers two commonly used normalization methods: Total Ion Current (TIC) normalization and Probabilistic Quotient Normalization (PQN). In the datasets presented in Figure 3, TIC normalization was applied, which standardizes each sample by dividing the intensity of each feature by the total ion current of the sample. In our analysis, TIC normalization revealed distinct clustering patterns between tissue and sample types in both datasets (Figure 3A and B), underscoring its utility in ensuring data comparability. TIC normalization is widely used in metabolomics due to its simplicity and effectiveness, particularly for datasets with consistent ionization efficiency across samples. Studies have demonstrated that TIC normalization enhances the reliability of comparative analyses by reducing technical variability while preserving biological variability [21,22]. ClusterApp also includes PQN, by calculating a median fold change between metabolites, which minimizes batch effects and corrects for technical variations during sample preparation or analysis. By using median fold changes relative to a reference, PQN improves data comparability and ensures reliable sample comparisons in metabolomic studies [23]. In ClusterApp users also have the flexibility to opt out of normalization entirely by selecting the “None” option, allowing them to analyze their data in its raw form if desired, or using a feature table normalized by other tool, as internal standard normalization, for example.

### Scaling

Data scaling, another key step in preprocessing, adjusts the range of features, ensuring comparability and enhancing the performance of multivariate data analysis. When features vary widely in scale, larger-scale features can dominate the analysis, leading to biased results. To address this, ClusterApp offers the most used scaling options in metabolomics field, pareto scaling, auto-scaling, and no scaling. Auto-scaling, which centers the data and scales each feature by its standard deviation, helps reduce the influence of high-variance features, ensuring a balanced contribution from all features in the analysis [15]. Pareto scaling is similar to auto-scaling but uses the square root of the standard deviation for scaling. This approach reduces the relative influence of large values while partially preserving the original data structure, making it particularly useful for emphasizing medium-variance features without distorting the dataset’s overall integrity [24,25]. In ClusterApp users also have the flexibility to opt out of scaling by selecting the “none” option, allowing them to analyze the data in its raw, unscaled form if desired, or using a feature table scaled by other tool.

As illustrated in Figure 3, the application of auto-scaling following normalization improved the clustering of key groups in both datasets. In the mouse tissue dataset (Figure 3A), combining auto-scaling with TIC normalization emphasized the separation between tissue types, unveiling distinct clustering patterns that were previously less noticeable. Similarly, in the coral life history stage dataset (Figure 3B), this approach enhanced the clarity of groupings corresponding to different life stages, effectively reducing within-group variability and sharpening distinctions between groups.

### Visualizing metabolomic variation with PCoA plots in ClusterApp

ClusterApp performs PCoA calculations and displays Emperor visualizations, which provides multiple tools to customize and enhance PCoA representations. For instance, users can distinguish groups with unique color codes and shapes, making it easier to identify and compare specific clusters within complex datasets. Figure 4 showcases PCoA plots generated using ClusterApp, illustrating metabolomic differences across sample types, genotypes, and phenotypes.

**Figure 4.**
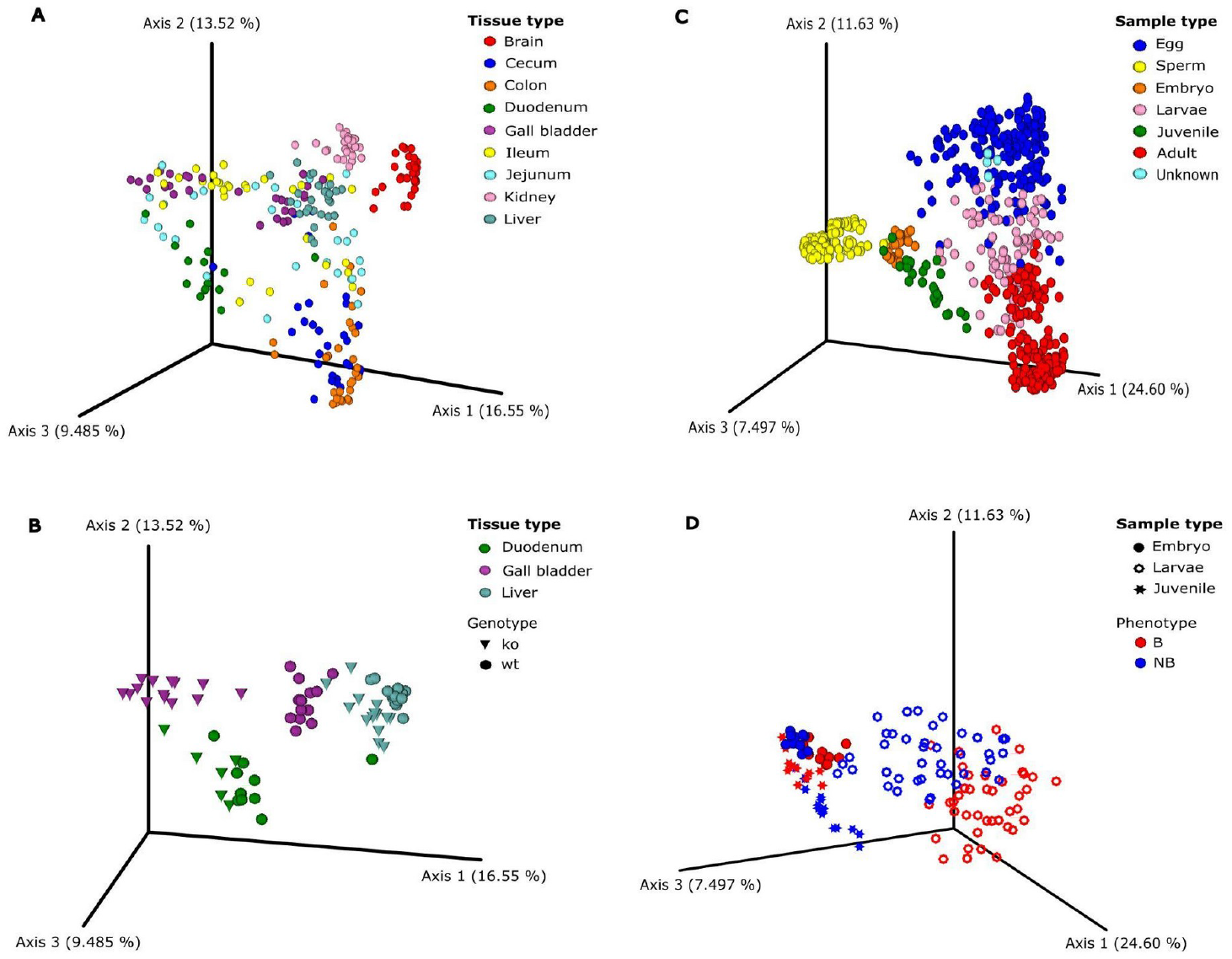
PCoA plots illustrating metabolome variation across different sample types, generated using ClusterApp. PCoA plots were produced with pairwise Bray-Curtis dissimilarity measures calculated for different A) tissue types, B) genotypes; BAAT^+/+^ (wt/wild type) and BAAT^-/-^ (ko/knockout) in mouse models, C) life history stages, and D) phenotypes; bleached (B) and non-bleached (NB) in coral samples. The plots are color-coded by tissue or sample type, with genotypes and phenotypes represented by different shapes.

In the mouse tissue dataset, the metabolomic variation between tissue types (Figure 4A) and genotypes (Figure 4B) was effectively highlighted. These plots were instrumental in evaluating the impact of the BAAT knockout (KO) on tissue metabolomic profiles, leveraging the app’s advanced visualization features to enhance data interpretation. Similarly, the distinct clustering patterns in coral datasets demonstrated differences across life stages (Figure 4C) and phenotypes, such as bleached (B) and non-bleached (NB) states (Figure 4D). Additional features, such as the ability to select or deselect groups (Figure 4B and 4D) and adjust node sizes, allow users to emphasize specific groups for clearer interpretation. These tools not only improve the clarity of data visualization but also enable researchers to focus on key elements relevant to their analyses, enhancing the overall utility of PCoA plots for metabolomic studies.

## Conclusion

The availability of easy-to-use tools for exploratory data analysis is essential to enable users without advanced bioinformatics knowledge to detect patterns in their data. The Web application ClusterApp fills a gap among the available tools performing data dimensionality reduction by providing a dynamic web server that allows useful data transformations coupled with PCoA calculation on multiple possible dissimilarity matrices. The tool can be accessed through a web interface or deployed locally using a docker image and used either by a local graphical interface or with Jupyter notebooks, enabling a wide range of use cases.

## Availability and requirements

**Project name:** ClusterApp

**Project home page:** http://ccbl-apps.fcfrp.usp.br/ClusterApp

**Operating system(s):** Platform independent

**Programming language:** Python, JavaScript

**Other requirements:** Docker

**License:** FreeBSD

**Any restrictions to use by non-academics:** e.g. licence needed

## Acknowledgments

We would like to acknowledge funding from the National Institutes of Health Grant R01AI145925 and R01DK140854 awarded to PI Quinn. We also acknowledge the supported by the São Paulo State Foundation (FAPESP), award number 2021/08235-3.

## References

1. Anderson M, Willis TJ. Canonical analysis of principal coordinates: a useful method of constrained ordination for ecology. Ecology. 2003;84(2):511–25.

2. Burges CJC. Geometric Methods for Feature Extraction and Dimensional Reduction - A Guided Tour. Data Min Knowl Discov Handb. 2009;53–82.

3. Jollife IT, Cadima J. Principal component analysis: a review and recent developments. Philos Trans R Soc A Math Phys Eng Sci. 2016 Apr 13;374(2065).

4. Wang M, Carver JJ, Phelan V V., Sanchez LM, Garg N, Peng Y, et al. Sharing and community curation of mass spectrometry data with Global Natural Products Social Molecular Networking. Nat Biotechnol 2016 348. 2016 Aug 9;34(8):828–37.

5. Tautenhahn R, Patti GJ, Rinehart D, Siuzdak G. XCMS online: A web-based platform to process untargeted metabolomic data. Anal Chem. 2012 Jun 5;84(11):5035–9.

6. Worley B, Powers R. Multivariate Analysis in Metabolomics. Curr Metabolomics. 2013 May 7;1(1):92–107.

7. Cambiaghi A, Ferrario M, Masseroli M. Analysis of metabolomic data: tools, current strategies and future challenges for omics data integration. Brief Bioinform. 2017 May 1;18(3):498–510.

8. Mildau K, Ehlers H, Meisenburg M, Del Pup E, Koetsier RA, Torres Ortega LR, et al. Effective data visualization strategies in untargeted metabolomics. Nat Prod Rep. 2024;

9. Pang Z, Zhou G, Ewald J, Chang L, Hacariz O, Basu N, et al. Using MetaboAnalyst 5.0 Part I: Optimizing parameters for LC-HRMS spectra processing. 2022 Jun 17;

10. Kuczynski J, Liu Z, Lozupone C, McDonald D, Fierer N, Knight R. Microbial community resemblance methods differ in their ability to detect biologically relevant patterns. Nat Methods. 2010 Oct;7(10):813–9.

11. Shlens J. A Tutorial on Principal Component Analysis. 2014 Apr 3;

12. Cox MAA, Cox TF. Multidimensional Scaling. Handb Data Vis. 2008;315–47.

13. Lozupone C, Knight R. UniFrac: a new phylogenetic method for comparing microbial communities. Appl Environ Microbiol. 2005 Dec;71(12):8228–35.

14. Hastie T, Tibshirani R, Friedman J. The Elements of Statistical Learning. 2009;

15. van den Berg RA, Hoefsloot HCJ, Westerhuis JA, Smilde AK, van der Werf MJ. Centering, scaling, and transformations: Improving the biological information content of metabolomics data. BMC Genomics. 2006 Jun 8;7(1):1–15.

16. Walach J, Filzmoser P, Hron K. Data Normalization and Scaling: Consequences for the Analysis in Omics Sciences. Compr Anal Chem. 2018 Jan 1;82:165–96.

17. Virtanen P, Gommers R, Oliphant TE, Haberland M, Reddy T, Cournapeau D, et al. SciPy 1.0: fundamental algorithms for scientific computing in Python. Nat Methods 2020 173. 2020 Feb 3;17(3):261–72.

18. Bolyen E, Rideout JR, Dillon MR, Bokulich NA, Abnet CC, Al-Ghalith GA, et al. Reproducible, interactive, scalable and extensible microbiome data science using QIIME 2. Nat Biotechnol. 2019 Aug 1;37(8):852–7.

19. Vázquez-Baeza Y, Pirrung M, Gonzalez A, Knight R. EMPeror: A tool for visualizing high-throughput microbial community data. Gigascience. 2013 Dec 1;2(1).

20. Dudzik D, Barbas-Bernardos C, García A, Barbas C. Quality assurance procedures for mass spectrometry untargeted metabolomics. a review. J Pharm Biomed Anal. 2018 Jan 5;147:149–73.

21. Dunn WB, Wilson ID, Nicholls AW, Broadhurst D. The Importance of Experimental Design And Qc Samples in Large-Scale And Ms-Driven Untargeted Metabolomic Studies of Humans. Bioanalysis. 2012 Sep;4(18):2249–64.

22. Broadhurst DI, Kell DB. Statistical strategies for avoiding false discoveries in metabolomics and related experiments. Metabolomics. 2006 Dec 28;2(4):171–96.

23. Dieterle F, Ross A, Schlotterbeck G, Senn H. Probabilistic quotient normalization as robust method to account for dilution of complex biological mixtures. Application in1H NMR metabonomics. Anal Chem. 2006 Jul 1;78(13):4281–90.

24. Eriksson L, Byrne T, Johansson E, Trygg J, Vikström C. Multi- and Megavariate Data Analysis Basic Principles and Applications. MKS Umetrics AB; 2013.

25. Gromski PS, Xu Y, Hollywood KA, Turner ML, Goodacre R. The influence of scaling metabolomics data on model classification accuracy. Metabolomics. 2015 Jun 1;11(3):684–95.

